# Highly Repeatable Tissue Proteomics for Kidney Transplant Pathology: Technical and Biological Validation of Protein Analysis using LC-MS/MS

**DOI:** 10.1101/2024.06.14.599091

**Authors:** Rianne Hofstraat, Kristina Marx, Renata Blatnik, Nike Claessen, Aleksandra Chojnacka, Hessel Peters-Sengers, Sandrine Florquin, Jesper Kers, Garry Corthals

## Abstract

Accurate pathological assessment of tissue samples is key for diagnosis and optimal treatment decisions. Traditional pathology techniques suffer from subjectivity resulting in inter-observer variability, and limitations in identifying subtle molecular changes. Omics approaches provide both molecular evidence and unbiased classification, which increases the quality and reliability of final tissue assessment. Here, we focus on mass spectrometry (MS)-based proteomics as a method to reveal biopsy tissue differences. For MS data to be useful, molecular information collected from formalin fixed paraffin embedding (FFPE) biopsy tissues needs to be consistent and quantitatively accurate and contain sufficient clinically relevant molecular information. Therefore, we developed an MS-based workflow and assessed the analytical repeatability on 36 kidney biopsies, ultimately analysing molecular differences and similarities of over 5000 proteins per biopsy. Additional 301 transplant biopsies were analysed to understand other physical parameters including effects of tissue size, standing time in autosampler, and the effect on clinical validation. MS data were acquired using Data-Independent Acquisition (DIA) which provides gigabytes of data per sample in the form of high proteome (and genome) representation, at exquisitely high quantitative accuracy. The FFPE-based method optimised here provides a coefficient of variation below 20%, analysing more than 5000 proteins per sample in parallel. We also observed that tissue thickness does affect the outcome of the data quality: 5 μm sections show more variation in the same sample than 10 μm sections. Notably, our data reveals an excellent agreement for the relative abundance of known protein biomarkers with kidney transplantation lesion scores used in clinical pathological diagnostics. The findings presented here demonstrate the ease, speed, and robustness of the MS-based method, where a wealth of molecular data from minute tissue sections can be used to assist and expand pathology, and possibly reduce the inter-observer variability.

## Introduction

In the field of medical diagnostics, accurately identifying and characterizing tissues is essential for patient care and effective treatment. Traditionally, tissue diagnosis has relied on histopathology, where tissue samples are examined under a microscope to detect structural and cellular abnormalities^1^. Traditional pathology techniques, while essential and widely used, suffer from subjectivity resulting in inter-observer variability, and limitations in identifying subtle molecular changes ^2^. As a result, there is a growing need for more objective and precise diagnostic tools to complement traditional methods. Molecular analysis of biopsy tissues offers a promising way to aid these shortcomings. However, comprehensive validation of reproducibility for specific applications is often lacking.

While histopathology remains a gold standard, advances in molecular techniques have revolutionized tissue assessment by enabling the analysis of proteins, the key functional molecules within cells and tissues. Proteins are crucial for all biological processes, governing cellular functions, signaling pathways, and gene expression. Their quantitative analysis, whether single proteins or full proteomes, mostly relies on liquid chromatography coupled to tandem mass spectrometry (LC-MS/MS). New MS-based workflows employing DIA approaches produce gigabytes of data in 10s of minutes, enabling fast identification and quantification of > 5000s of proteins at exquisite sensitivity and specificity, through the correlation of peptide fragmentation patterns with human genome information^3^. This untargeted view on the proteome content of tissues enables comparison of protein expression patterns across different tissues or disease stages, thereby aiding in disease classification, prognostication and support patient tailored treatment decisions^4,5^.

In the current study we use the newest generation of LC integrated MS-based proteomics technology (EvosepOne coupled with a timsTOF HT) to gain insight into the diagnostic accuracy and workflow reproducibility for relative quantification of proteins in FFPE patient tissues. First, we evaluated the consistency and robustness of the method by optimizing and validating our integrated FFPE-tissue workflow on a subset of 36 samples, ranging in biopsy size and visual signs of inflammation. In the second stage, evaluation of 301 kidney biopsies from the DEEPGRAFT digital pathology cohort^6^ were used to understand if practical aspects such as tissue size and LC-MS integration influenced assessment of clinical rejection indicators. Overall, the current setup, EvosepOne coupled with a timsTOF HT operating in data-independent acquisition mode with parallel accumulation and serial fragmentation (dia-PASEF mode), provides broad proteome coverage with high sensitivity and quantitative accuracy, enabling the detection of biologically relevant differences in a fraction of the time required by other methods^7^.

## Materials & Methods

### Sample Selection

For this study we used 2 separate sample collections: 1) kidney biopsy tissues for the assessment of repeatability, 2) kidney biopsies from a large transplant cohort for the assessment of clinical rejection indicators. The tissue selected for the assessment of the robustness and accuracy of the instrument and sample preparation was based on the size of the biopsy and large visual signs of inflammation. The block needed to have enough tissue to obtain 6 samples of 5 um thickness and 6 samples of 10 um thickness. The inflammation in the validation samples was required to resemble the transplant samples and test if immune related proteins were detected. The transplant cohort consist of kidney transplant samples comprising all patients between January 1, 2000 and June 1, 2018, who had kidney transplant biopsy performed at the Amsterdam UMC, location AMC, and which were included in the initial screening of the DEEPGRAFT cohort^6^. There were no exclusion criteria other than the availability of the biopsy. From the 1048 kidney transplant biopsies a selection of 301 biopsies was made for this study, which contained common diagnoses occurring in kidney transplant complications (Table 1). All kidney transplant biopsies were retrospectively re-evaluated according to the 2019 Banff classification by an expert kidney pathologist (SF)^8^. Experiments were performed in accordance with the Declaration of Helsinki and were approved by the local ethics and privacy committee at Amsterdam UMC, waiving informed consent (number 19.260).

**Table 1.**
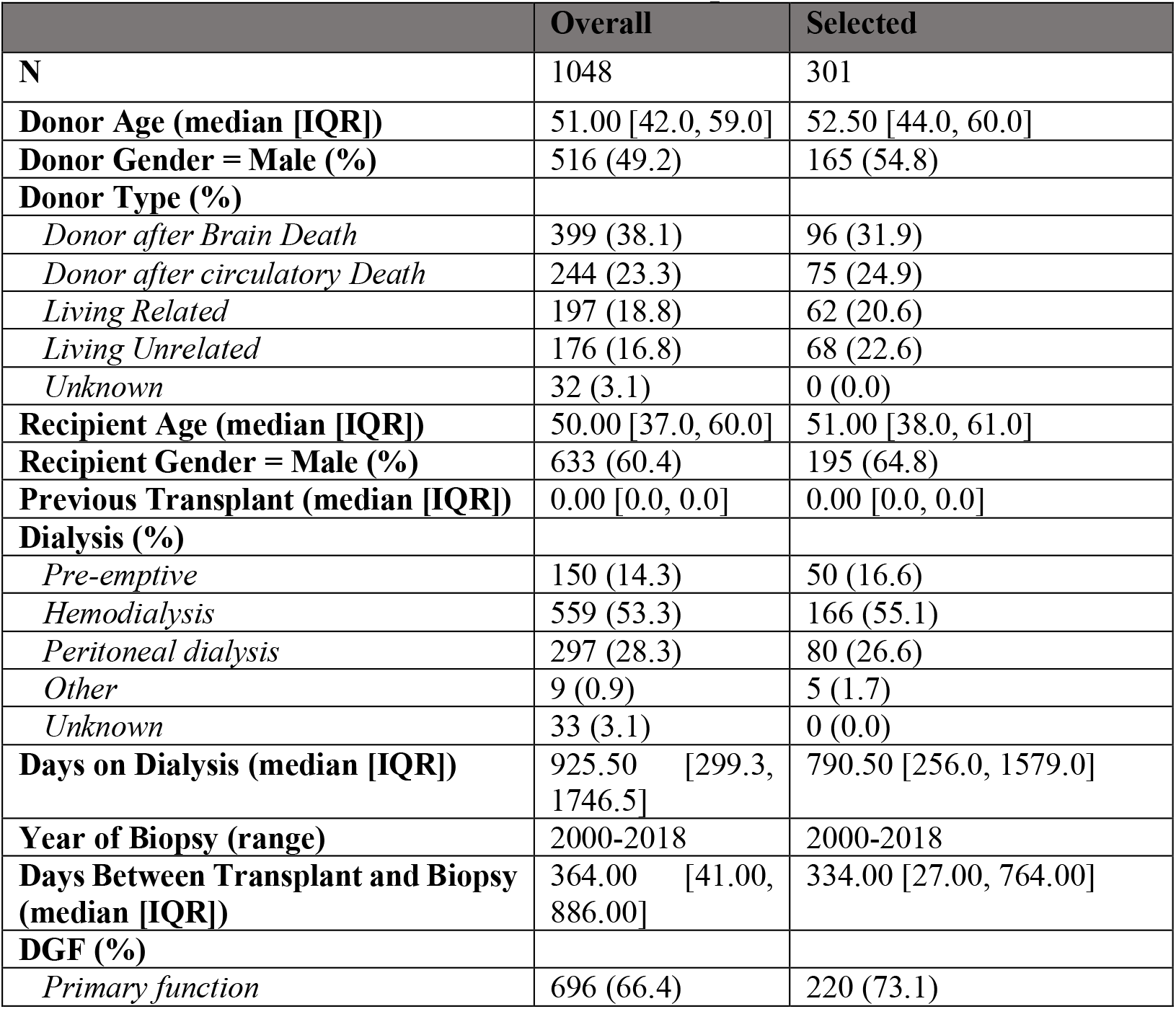
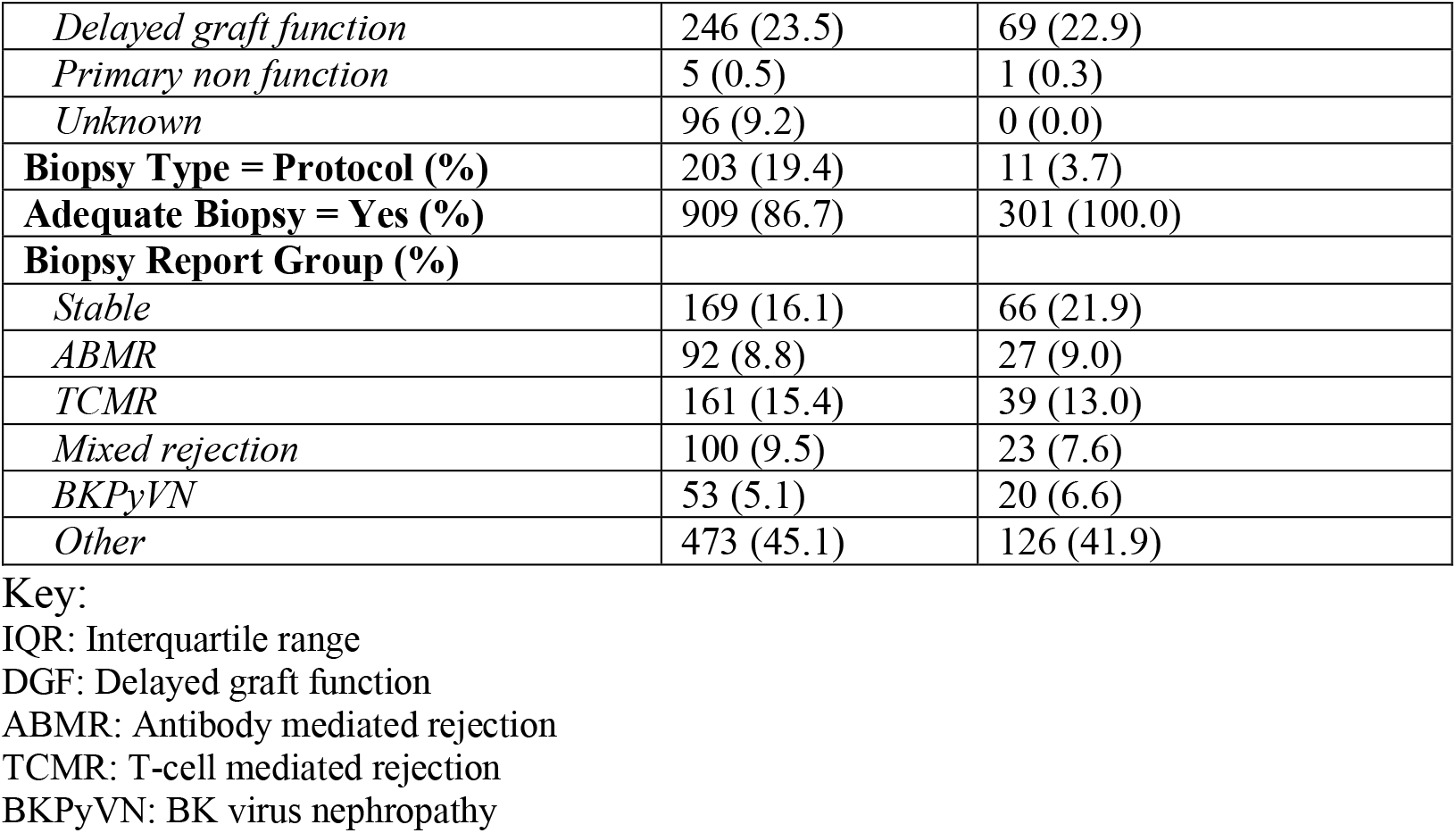
Baseline characteristics of DEEPGRAFT and proteomics cohort.

### Kidney biopsy processing

FFPE kidney transplant biopsies were fixated overnight in 4% buffered formalin, the next day embedded in paraffin and subsequently stored at room temperature until analysis. Tissue samples were generated with a standard clinical pathology lab microtome at 5 and 10 μm thickness. The samples were cut very slowly to create a rolled-up piece of tissue which could be collected directly into an Eppendorf tube without the need for transfer to a glass slide first. This significantly simplified the tissue collection and processing, as removing material from glass slides often generates static electricity, increasing the risk of tissue loss during the process. All subsequent processing is done within the same Eppendorf tube.

### Materials and reagents

Water, Acetonitrile (ACN), Formic acid (FA), Trifluoracetic acid (TFA), Ammonium bicarbonate (NH4HCO3), Iodoacetamide (IAA), Dithiothreitol (DTT), Trypsin and C18 SPE cartridges were obtained from Merck (KGaA, Darmstadt, Germany). The EV1109 column was obtained from Evosep (Evosep Biosystems, Odense, Denmark).

### Biopsy preparation for LC-MS/MS

FFPE tissue preparation was based on an originally reported method by Dapic et al^9^. FFPE tissues were first deparaffinized by incubating the tissue in the Eppendorf tube in a thermomixer at 40℃ and 850 revolutions per minute (rpm) and adding xylene for 10 minutes (min) to remove the paraffin, we repeated this 4 times^9^. Next, the tissue was incubated in the thermomixer at 60℃ and 850 rpm for 4 min in ethanol, a process that was repeated three times. Finally, we incubated the Eppendorf tube with an open lid in the thermomixer at 95℃ to evaporate the remaining ethanol. Samples were incubated in 100 μL of 30% acetonitrile v/v 100 mM ammonium bicarbonate buffer for 90 minutes at 95 °C in the thermomixer at 1050 rpm. After the initial antigen retrieval phase, an extraction and a reduction solution were prepared for each sample by dissolving 72 mg of urea in 100 µL of a 30% acetonitrile (ACN) solution in 100 millimolar (mM) ammonium bicarbonate. Subsequently, 100 µL of this prepared solution was added to each sample, followed by a further incubation period of 30 minutes at 37 °C in an oven. Hereafter, 2.5 μL DTT (700 mM) was added to each sample and incubated for 30 min at 37 °C in the thermomixer. Further, 9.2 μL IAA (700 mM) was added to the samples and incubated for 30 min at 37 °C. Finally, samples were diluted with 120 μL of 1 molar (M) ammonium bicarbonate and 880 μL of MS grade water (H_2_O). To each sample, 5 ng of trypsin was added per mm^2^ of tissue and was incubated overnight at 37 °C in the thermomixer. The next morning another 5 ng of trypsin per mm^2^ was added and incubated at 37 °C for 4 hours. After digestion, samples were purified on a SPE cartridge, freeze-dried and stored at −80 °C until instrumental analysis.

### MicroLC-ESI-MS/MS

The samples were reconstituded using 24 µL of 2% acitonitrile in 98% water and 0.1% of TFA, 1 µL was taken from this and diluted using 59 µL of 0.1% formic acid in water. 20 µL from each sample was deposited onto Evosep tips and inserted in the Evosep tray. The samples were analyzed using an Evosep One: EV-1000 (Evosep, Odense, Denmark) with the Evosep 60SPD method ^10^. The 60SPD method has a 21-minute gradient and a cycle time of 24 minutes where sample selection was performed in a randomised order. The analytical column was equilibrated at 2 μL/min. The gradient flow is 1 μL/min and increased to 2 μL/min for washing. The peptides were eluted with the mobile phase gradient starting at 0% mobile phase B, which gradually increases to 40% in 20 minutes, the last minute of run time the mobile phase B increases to 90% followed by 2 minutes of washing at 90% mobile phase B. Mobile phase A was composed of formic acid in water, and mobile phase B of 0.1% formic acid in acetonitrile. Detection of the peptides was performed on a timsTOF HT (Bruker, Bremen, Germany) in dia-PASEF mode, with a mass range of 100-1700 Da, ion mobility range of 0.85-1.3 V*s/cm^2^, cycle time of 0.95 s and a ramp time of 100 ms.

### Repeatability assessment

Using repeated measurements, we compared the effects of biopsy sample thickness on the total protein yield, to determine the optimum size for consistent results. Furthermore, the sample handling was assessed by evaluating 6 biological identical samples to determine total protein yield. In this paper, the six biological samples are referred to as the biological replicates, and the three repeated injections performed on each of these samples are termed the technical replicates. The duration was carefully examined by assessing potential degradation or changes in protein yield over time, thereby determining a time frame that ensures stability. We kept the conditions of the experiments constant, including the same measurement procedure, same operator, same measuring system, and same location.

### Data Analysis

Raw dia-PASEF files were processed using DIA-NN^11^, predominantly in default setting (version 1.8) without spectral library. Instead, the human Uniprot Reference Proteome without Isoforms FASTA file was imported (April 2023 release with 26,751 protein sequences). The protease we used was Trypsin-P, allowed for 1 missed cleavage and cysteine carbamidomethylation and N-terminal methionine excision were selected as modifications. We specified a minimum of 7 and maximum length of 30 amino acids for peptides, the precursor charge range was between 1-4. The precursor m/z range was between 299-1250 and the fragmentation ion m/z range between 150-1650. The precursor FDR was set to 10% and protein detection at 1%. For the algorithm section, we deselected Unrelated runs and match between runs (MBR), and left the mass accuracy, MS1 accuracy and scan window at 0. The neural network classifier was set at double-pass mode, for protein inference we used protein names (from FASTA), we set quantification strategy to robust LC (High accuracy), cross-run normalization on global, library generation on smart profiling and speed and RAM usage was set to optimal results.

Two normalization procedures were conducted. The first normalization is performed in the data processing software DIA-NN, using retention time-dependent normalization on 5 μm and 10 μm samples separately^11^. This normalization corrects for undesired technical effects, which leads to the depletion of hydrophobic or hydrophilic peptides and potentially, at the same time corrects for issues caused by e.g. spray instability. DIA-NN also performs a MAX-LFQ normalization^12^. Count normalization was performed on both 5 μm and 10 μm together using mean normalization with the DESeq2 software package (version 1.40.2) in R^13^.

The data analysis was implemented through R (version 4.3.1). First the data was imported using the diann package (version 3.6.2). To determine the amount of variation differences between 2 sample groups a principal component analysis (PCA) was performed using the DeSeq2 (version 1.40.2) package. This package was also used for the biological visualization through a heatmap and volcano plot. The differences in variation between 2 groups of samples was visualized using the tidyverse package containing ggplot2 (version 3.3.2), creating violin and density plots. Furthermore, the significance was assessed through the Kruskal–Wallis or Mann–Whitney test depending on two or more groups, providing us with the p-value. A p-value < 0.05 defined significance. The ggplot2 package was used to create a scatter plot that includes a regression line and an R-squared value, to determine how much of the data is represented by the regression line.

## Results

### DEEPGRAFT cohort description

We included 301 patients of the kidney transplanted patients with a kidney biopsy taken between the year 2000 and 2018 in this study. Table 1 shows the characteristics of the patients that were enrolled in the study. Mean donor age was 52.5 years (standard deviation, SD 16.0), and recipient age was 51.0 years (SD 16.8). Majority of patients received a living donor kidney (43.2%), followed by brain-death (31.9%) and circulatory-death (24.9%). Indications for biopsy included most of the patients suffered from rejection (29.6%), comprising a combination of 13.0% T-cell mediated rejection (TCMR), 9.0% antibody mediated rejection (ABMR) and 7.6% mixed rejection. Other patient groups had a stable transplant (21.9%), followed by BK virus nephropathy (BKPyVN, 6.6%). The remaining biopsy group (41.9%) is comprised of small other categories.

### Technical Repeatability

To assess the instrument’s repeatability, we consecutively injected the same biological sample three times. There was substantial overlap among the 10 μm samples indicating a high level of consistency in the measurement, visualized through a PCA plot (Figure 1A). In contrast, the 5 μm samples exhibited a broader dispersion, even though the 2^nd^ principal component 2 (PC2) variation remains low. Upon further examination of the PCA for each group separately, the 10 μm biopsy shows a variation of 22% in PC1 and 16% in PC2. In contrast, for the 5 μm samples, PC1 accounted for 32% and PC2 for 18% of the variation, indicating more variability than the 10 μm samples. To further scrutinize the variation among the same three replicates of six samples, we visualized the coefficient of variation (CV) of proteins with a violin plot (Figure 1A, and B). This plot illustrates minimal variation within the 10 μm samples, while the 5 μm samples exhibit more pronounced diversity. The interquartile range (IQR) of coefficient of variation of the 5 μm samples is between 15.5% and 27.2%, which is much larger than the mean of the 10 μm samples, which is between 9.5% and 11.4% (Supplimentary Table 1). The 10 μm samples have only one protein with a CV higher than 20% at the 75^th^ percentile (20.2%).

**Figure 1.**
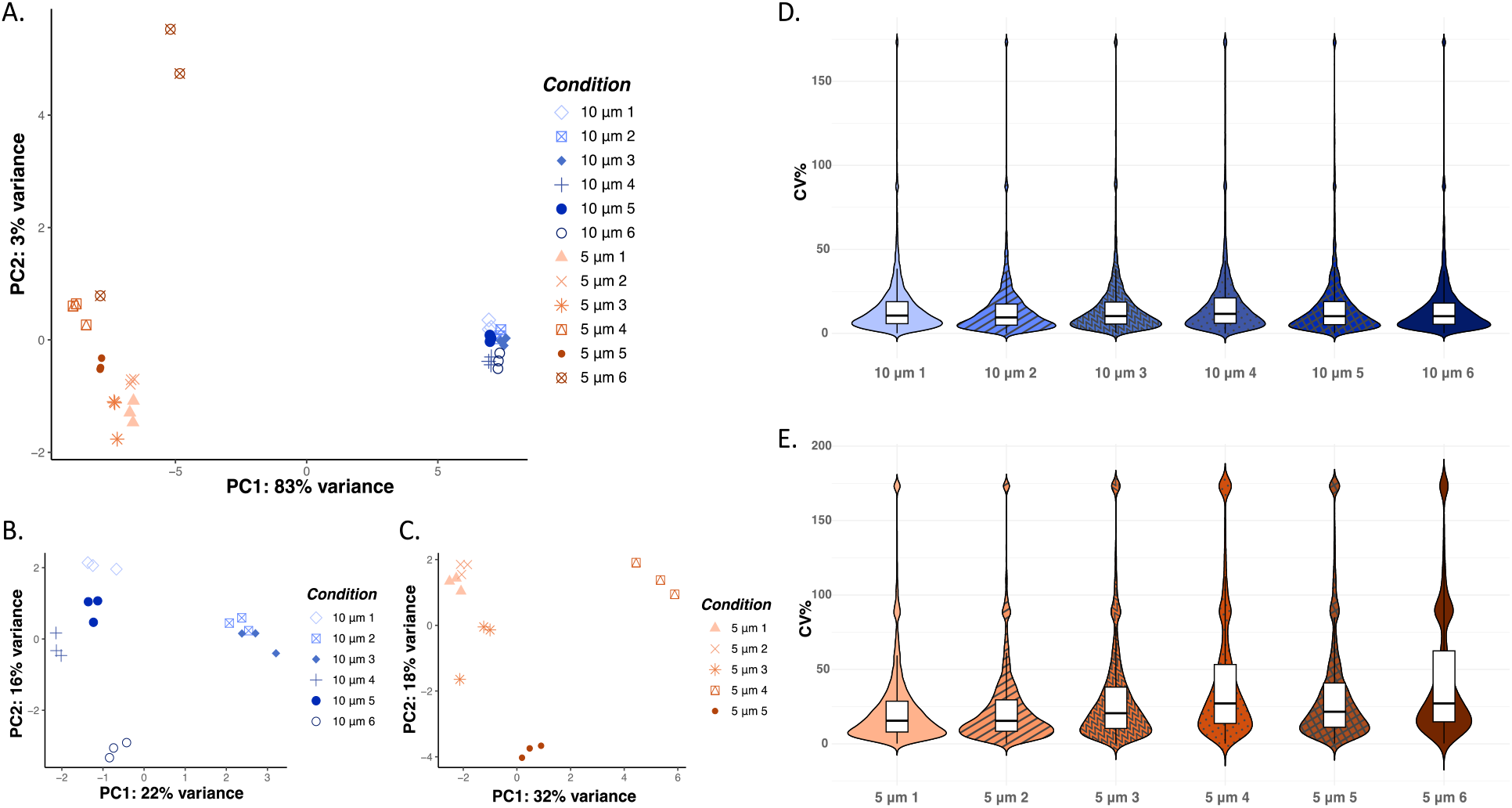
Assessment of technical repeatability. A. Principal component analysis (PCA) of 5 μm and 10 μm samples, each containing 6 identical samples which were injected into the system 3 times, showing the largest difference between the 2 groups and clustering within the groups. PC 2 shows the presence of two outliers due to the instrument stopping during the runs (still only accountable for 3% of the variation in the data). B. PCA of 10 μm thick biopsies. C. PCA of 5 μm thick biopsies. D. Coefficient of variance (CV) of 10 μm biopsy replicates containing all proteins present even when it was only present in one of the replicates resulting in percentages above 100%, visualized using violin plot containing a boxplot. E. Violin plot combined with boxplot of the CV of 5 μm replicates containing all proteins present even when it was only present in one of the replicates resulting in percentages above 100%.

### Biological Repeatability

To determine the consistency of sample preparation, we injected different samples of the same tissue block (biological replicates). As depicted in Figure 2, the PCA for biological samples also shows the largest variation is between the 2 different thicknesses. To determine the variation, a PCA was performed of each group separately. When examining the plots separately we observed that there was still more variation between the 5 μm samples with a PC1 of 49% and a PC2 of 26%. In comparison, the 10 μm samples had 34% of variation in PC1 and 28% in PC2. Further investigation via a combined violin and box plot revealed that the median CV of all proteins for the 5 μm thickness group was 19.4%, while the median CV for the 10 μm thickness group was 10.2% (Figure 2 B and C). It is worth noting that these findings contain all the proteins detected in all the samples, as is evident at the upper portion of the violin plot, which also highlights specific proteins unique to individual samples. When inspecting the IQR for the 10 μm replicates, it becomes apparent that, the 75^th^ percentile of the detected proteins is at 15.4% CV, in contrast, the 5 μm thickness group exhibits 30.7% CV at the 75^th^ percentile (Supplementary Table 1).

**Figure 2.**
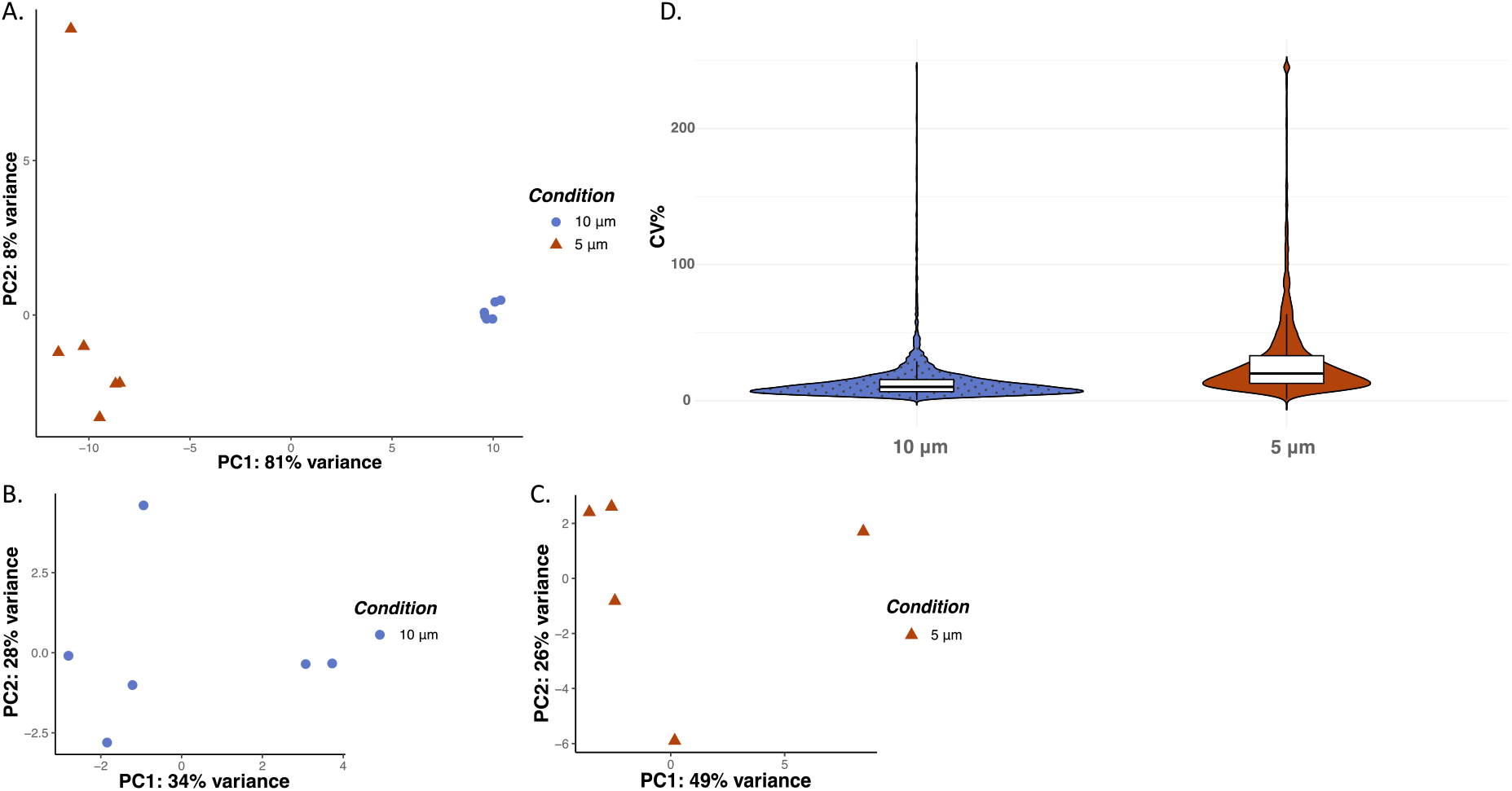
Assessment of biological repeatability (A-D). A. Principal component analysis (PCA) of 5 μm and 10 μm samples, each containing 6 identical samples using the mean of the technical replicates, showing the largest difference between the 2 different groups. PC 2 shows the presence of one outlier due to stopping of the instrument during the run, however, is still only responsible for 8% of the variation in the data. B. PCA of 10 μm thick biopsies. C. PCA of 5 μm thick biopsies. D. Coefficient of variance of 10 and 5 μm replicates visualized using violin plot containing a boxplot, containing all proteins present even when it was only present in one of the replicates resulting in percentages above 100%.

### Biological Differences between Sample Thickness

Next we determined the difference in biological information between 10 μm and 5 μm thickness. Figure 3A shows that there is a group of proteins only present in 10 μm or in 5 μm. We observed significant differences in protein yield between 10 μm (5570 proteins) and 5 μm (4793 proteins) (visualized through a volcano plot in Figure 3B). The difference in proteins abundance between both sample sets were not related to the immune system. A gene set enrichment analysis confirmed that immune related pathways were not different between 10 µm and 5 µm. Also Figure 3C shows that there are no pathways significantly up or downregulated.

**Figure 3.**
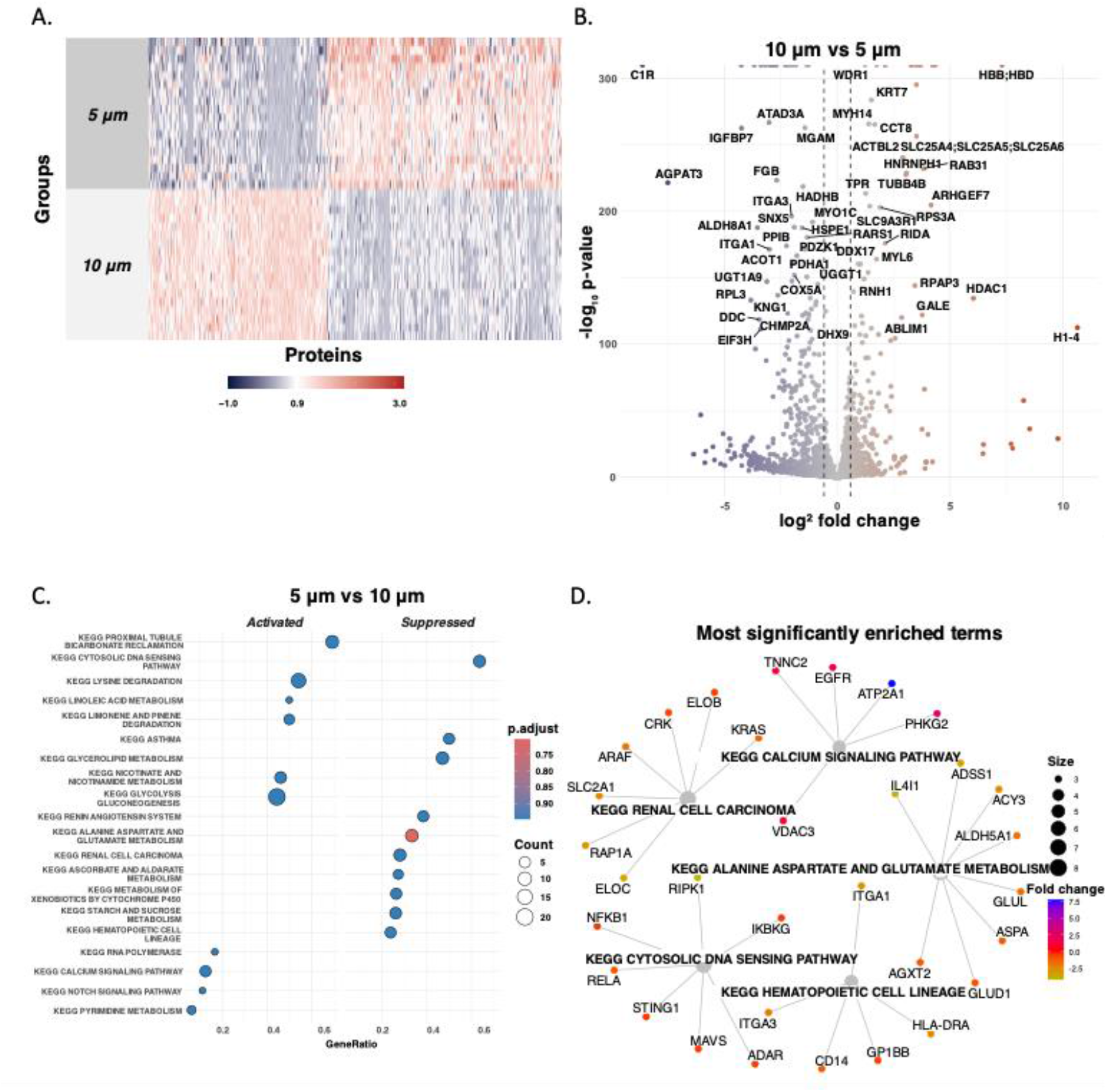
Assessment of the molecular differences between 5 μm and 10 μm repeats. A. Heatmap of the 2 groups, where blue suggests down regulation, white represents no variation, and red shows a higher expression between the two groups. B. Volcano plot of up and down regulated genes correlated to proteins detected with LC-MS/MS. C. Gene set enrichment analysis visualized using a dotplot, showing activated and suppressed pathways in 5 μm relative to 10 μm. D. Overview of the most significantly enriched pathways in 5 μm relative to 10 μm.

### Effect of sample characteristics and handling on protein quantity

Following assessment of repeatability in sample preparation and analysis, it is essential to verify if the analysis does not affect large cohorts of clinical samples, therefore, we leveraged a larger cohort of 301 transplant biopsies to investigate this. This included evaluating the impact of biopsy size, number of glomeruli, type of kidney rejection, lesions scores, batch effects, as well as degree of inflammation and fibrosis.

### Batch effect

The samples underwent a continuous injection into the LC-MS/MS, which inevitably resulted in varying durations of exposure to the autosampler (0-2 days). Notably, the Evosep system used, lacks a temperature-controlled compartment — a feature commonly found in most LC systems. Sample selection was performed in a randomised order using the Evosep system. To assess the potential impact of this non-temperature-controlled environment on our samples, we explored the number of proteins detected in each sample in chronological order. Figure 4 illustrates no visible trends (F_1,302_ 0.9963, P = 0.319), suggesting no decline in protein detection over time or any indication of distinct batch effects.

**Figure 4.**
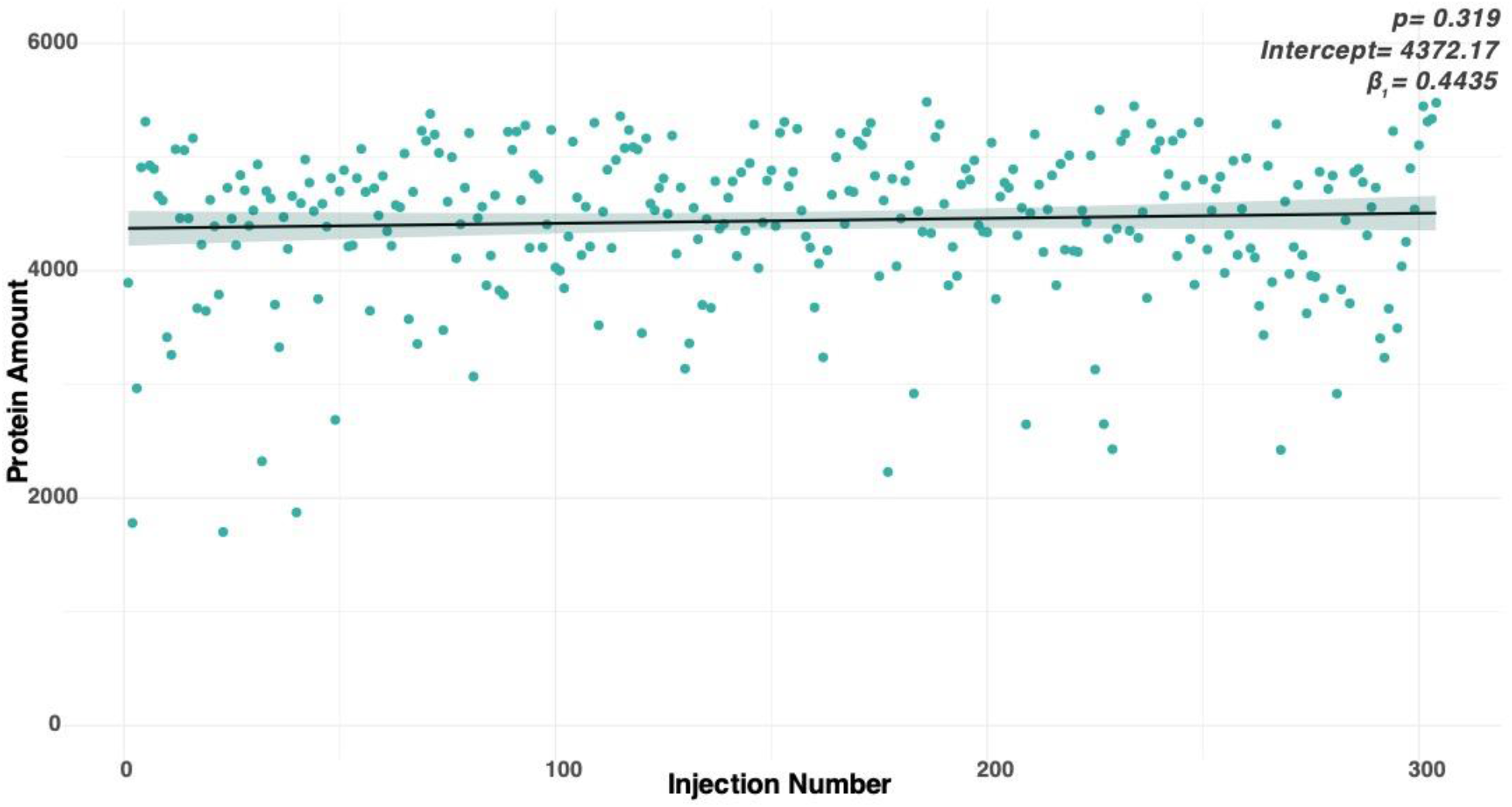
Number of proteins detected depending on when the sample was injected.

### Size

Within our transplantation cohort of 301 samples, we encounter a diverse array of biopsy areas. Despite consistently using the same thickness for each sample, this diversity in biopsy size might impact the total amount of tissue utilized. To investigate the influence of sample size on our analysis, we examined the amount of protein present in samples of multiple size degrees. The histogram in Figure 5A illustrates that smaller samples typically have a lower number of proteins detected, whereas larger samples generally contain a greater number of proteins. On average, the group of the smallest biopsies had 540 less proteins than the group of the largest biopsies (H[2]=22.154, P =1.547e-05, Figure 5A). What also became evident was the increased variability in the number of proteins within smaller samples compared to the larger samples (Figure 5B).

**Figure 5.**
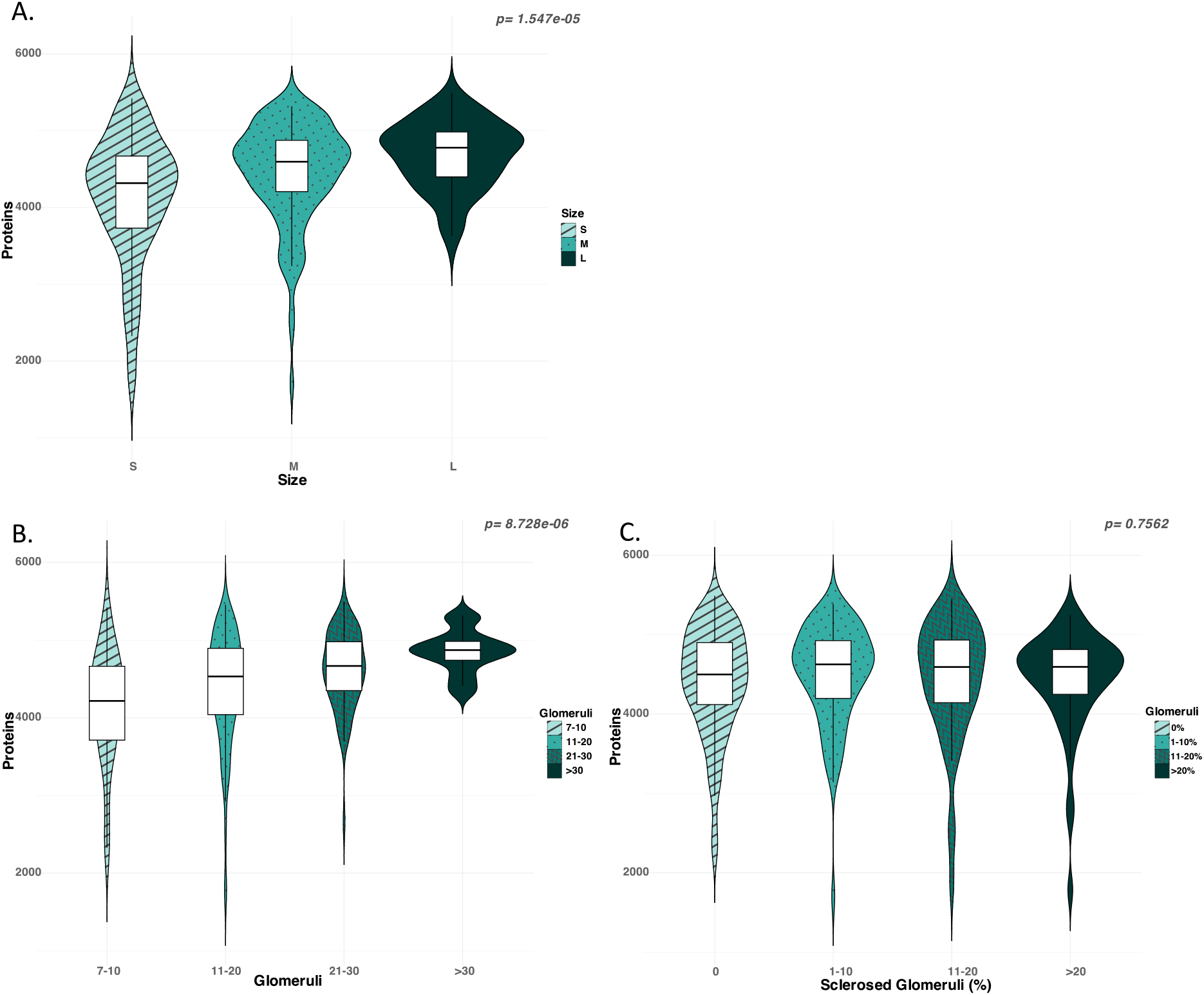
Influence of the size of the sample on the yield of number of proteins. A. Estimated size of proteins correlated to the number of proteins which were detected. B. Number of glomeruli which correlates with the amount of cortex found in the biopsy. C. Percentage of sclerosed glomeruli, which indicates the amount of damage in the kidney. No significant differences were observed between groups.

A typical kidney biopsy has a variable size of cortical and medullar sampling, but diagnostic assessment is typically done on cortex alone (in the case of the Banff classification for allograft pathology)^8^. We used the number of glomeruli as a proxy for cortical sampling. Figure 5B shows that the protein yield was different across groups with different amount of glomeruli present (0-10, 11-20, 21-30 and >30, H[3]=26.184, P =8.728e-06). Additionally, we delved into the correlation between sclerosed glomeruli and the analyzed proteins, to determine whether a large number of proteins that we detect comes from the glomeruli (Figure 5C). This shows no difference in protein yield in the different groups of sclerosed glomeruli (0, 10-20, 21-30% and >30, H[3]=1.1866, P =0.7562).

### Diagnostic group & Lesion Scores samples

The Banff classifications serve as the gold standard for diagnosing kidney transplant pathology^8^. This classification system includes various lesion scores, which, when combined, provides a specific diagnosis. It is imperative that the distribution of proteins among these different groups remains similar to eliminate bias when analyzing molecular information. Given the diversity of lesion scores within our cohort, we tried to assess whether there is a consistent distribution across all scores. However, as depicted in Figure 6, notable differences occurred within a single lesion score. This is particularly evident in the tubulointerstitial inflammation through Interstitial inflammation (i) and tubulitis (t) lesions (it-score, Figure 6D), where for different scores different protein yields were observed, however, not significant (H[3] = 3.7043, P= 0.2952). Next, we examined the protein yield in biopsies across different diagnostic groups (stable vs rejection vs other, Figure 6E). In contrast to the lesion scores, Figure 6A-D reveals an overall similarity in protein yield across the various diagnostic categories (H[2] = 0.13401, P= 0.9352).

**Figure 6.**
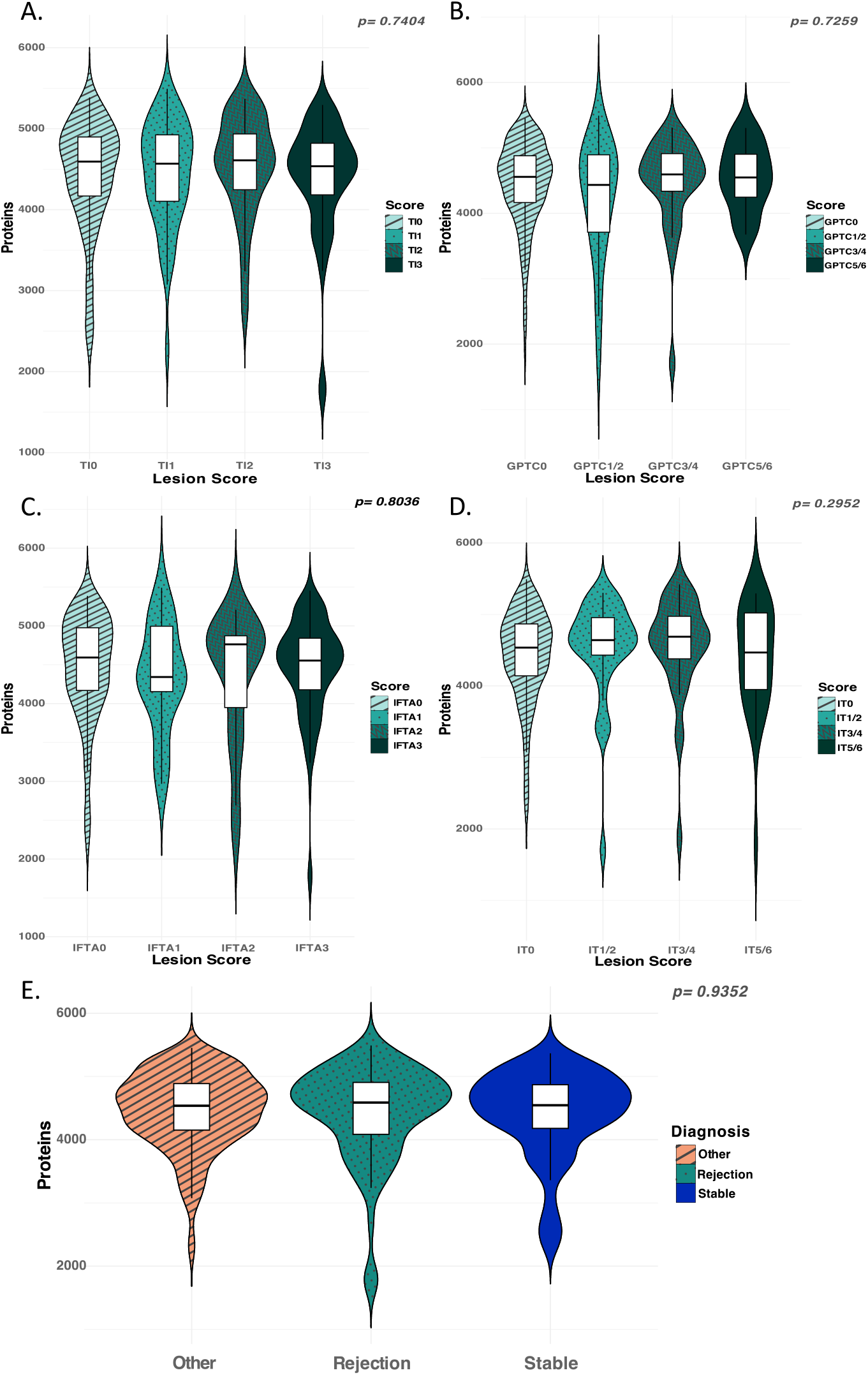
Assessment of bias due to sample selection. A. Number of proteins per total inflammation (TI) lesion scores. B. Number of proteins in microvascular inflammation through Glomerulitis (g) and peritubuliar capillaritis (pct) lesion scores. C. Number of proteins correlated to interstitial fibrosis and tubular atrophy using the IFTA lesion score. D. Number of proteins in tubulointerstitial inflammation through Interstitial inflammation (i) and tubulitis (t) lesion scores E. Number of proteins per diagnostic group to determine if any biological differences are not due to the measurement.

### Targeted assessment of lesion-specific proteins as positive control

To validate our analysis, we examined two markers known to be responsible for the pathological processes of inflammation (leukocyte common antigen, CD45) and fibrosis (collagen type 1, COL1). In transplantation, but also in native kidney diseases, the assessment of disease activity and chronicity holds significant clinical importance, influencing treatment decisions—whether to initiate, modify, or cease treatment. Activity can be determined by the burden of total cortical inflammation (the Banff ti-score)^14^. CD45 (or leukocyte common antigen; LCA), a protein expressed on all leukocytes, and correlates positively with an increase in ti-score (H[3]=123.79, P= 2.2e-16) (Figure 7A). Chronic damage is determined by the burden of fibrosis (scarring), mostly of the tubulointerstitial compartment. Collagen type 1 exhibits a positive trend correlated to the increase in interstitial fibrosis and tubular atrophy (H[3]= 23.044, P= 3.953e-05, Banff IFTA score, Figure 7B^15^).

**Figure 7.**
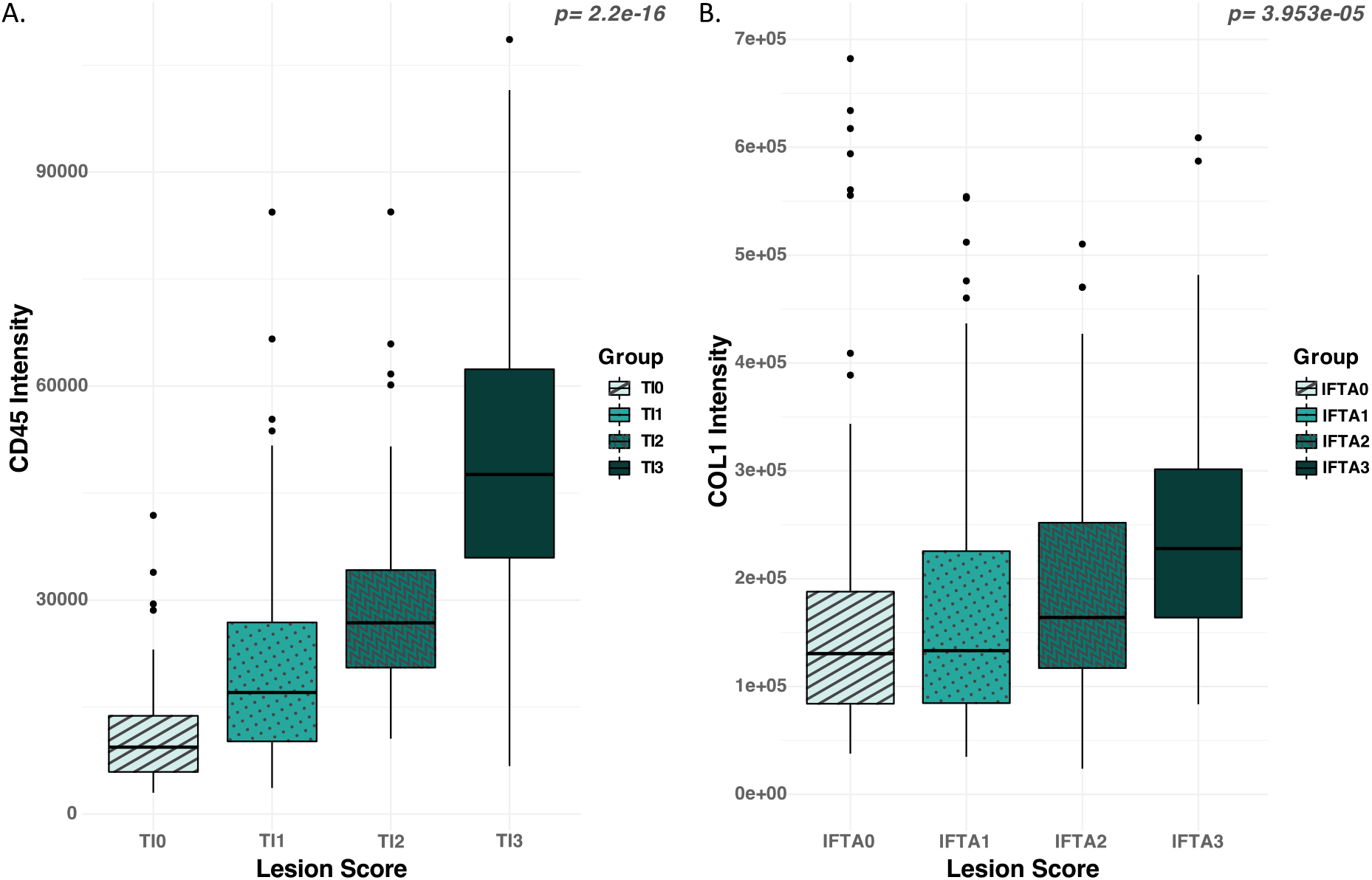
High-throughput quantitative proteomics captures the expected correlation between (A) the degree of leukocyte infiltration as measured by CD45 (Leukocyte Common Antigen) and determined by histologic scoring (Banff total inflammation [ti] score) and (B) the amount of collagen present through COL1 intensity using IFTA histological scoring (Banff interstitial fibrosis and tubular atrophy score)

## Discussion

Kidney transplantation has a 70-year history, but challenges remain in achieving optimal long-term outcomes for patients^16^. Some shortcomings can partially be attributed to significant interobserver variability among pathologists, a challenge that is intensified by the inherent complexity of diseased kidney transplant tissue^17^. Even with explicit guidelines provided to pathologists, substantial variability persists^18^. Incorporating data-driven and unbiased methods may reduce variability and provide a more accurate assessment of the underlying environmental or biological processes that play a role in post-transplant disease states. The application of proteome-based methods to tissue analysis revealing huge datasets of proteins for a given patient sample and observing unambiguous specificity and quantification with exquisite sensitivity, offers great potential to pathology. Importantly, such MS-based methods offer specific information on proteins in an unbiased way. Besides these advantages, proteins represent the functional output of gene expression governing and catalysing all molecular processes, including key molecules in the immune system, and structural layout in tissues. Thus, the analysis of the tissue-specific proteome per patient sample represents a tool to reduce variability and detect molecular changes before they become visually apparent in the form of pathological lesions, thus potentially contributing to an earlier and more accurate diagnostic process^19^.

In this study we successfully validated a integrated workflow consisting of sample preparation, from 5 and 10 µm FFPE tissues, with a rapid (60 samples per day, SPD) and comprehensive MS-based method capable of quantifying over 5000 proteins per tissue sample. We primarily targeted two aspects of the overall workflow: Firstly, the repeatability of the MS-workflow, and secondly, we investigated the clinical relevance of the information obtained. Both are of primary importance in evaluating suitability for the clinic.

Three critical findings are: 1) the LC-timsTOF HT, operated at high throughput (60 SPD) in dia-PASEF-mode, has acceptable technical and biological repeatability (majority of all proteins <20% CV); 2) measurements are stable over multiple runs through the study; and 3), which is of clinical relevance the protein content (not the protein yield) is reflective of the severity of histological lesions known to be important for kidney transplantation. Although not specifically tested, these findings can be directly translated to biopsies from native kidney diseases since the collection and processing of material is equal.

In general, there is only little literature dealing with the FFPE proteomics of kidneys. We and others have previously reported methods for the feasibility of analysis, and primarily explore the distinctions between Fresh Frozen (FF) and FFPE tissue, with limited emphasis on clinical utility or relevance ^9,20–22^. Here, the primary objective was to assess the viability of proteomics as a potential diagnostic tool in pathology, resulting in the sole use of FFPE material, which is in line with diagnostics and easier storage.

The technical feasibility of the method presented here is demonstrated by the minimal changes in protein composition observed across repeated measurements. It is important to underscore a difference between this method to previously published methods^9,20–22^. In previous tissue proteomics studies, samples are typically mounted on slides, measured for size, and specific areas are selected for analysis. However, our experience teaches us that scraping samples from slides generates static electricity, hampering the sample preparation and increasing the risk of (at times complete) sample loss. To address this issue, we devised a method where tissue is rolled directly from the microtome and placed into an Eppendorf tube. While this approach eliminates the possibility of precisely measuring area, it does not result in the loss of protein quantity but does result in varying tissue sizes between samples. With this knowledge we investigated the impact of size difference on protein identification, demonstrating less variation of the number of detected proteins in larger samples compared to smaller ones. Importantly, in our large cohort of 301 samples, we found no association between sample size and diagnosis, indicating that tissue size does not significantly affect diagnostic outcomes and does not cause bias.

Considering the elimination of mounted samples, which bypasses concentration measurement, we measured two commonly used thicknesses of 5 µm and 10 µm. The MS-data presented here shows that one can achieve reproducible measurements of individual samples as small as 5 μm. However, as tissue thickness increases, we observe a significant reduction in technical variation. Ten µm thickness provides a greater repeatability, where almost all (>) 5000 proteins exhibit a CV below 20%, which is acceptable given the complexity of our samples^23^.

Acute and chronic rejection have a large contribution in allograft kidney loss and can be seen as a huge factor limiting long term graft survival^24^. Acute rejection results in a rapid decline of the kidney function and is characterized by inflammatory burden in each of the renal anatomic compartments^25^, summarized into the Banff total inflammation (ti)-score. On the other hand, chronic rejection typically shows a slower and more progressive decline of the allograft function, which is considered irreversible and characterized by the burden of extracellular matrix accumulation in the various renal compartments^26^, summarized into the Banff interstitial fibrosis and tubular atrophy (IFTA)-score. On the molecular level, these lesions are represented by the total leukocyte influx (CD45^27^) and deposition of collagen type 1 (COL1). We determined that if the pathologist sees an increase of these lesions, we also see an increase of the correlated protein in our analysis. This further shows that the analysis also provides us with the expected biological information, however, in an objective and quantitative form.

While we are enthusiastic about these new findings, it is important to acknowledge limitations of this study. While many aspects of LC-MS/MS robustness were tested, certain parameters remain unexplored. Assessing instrument and sample robustness would require analyzing the same sample on different days and revealing degradation trends over time. Also, reproducibility, another critical facet, could be assessed by preparing and analyzing the sample by different laboratories. These are important parameters to be assessed in future studies.

## Conclusion

Primarily, the study shows the robustness of tissue proteomics for FFPE samples using roll-ups of varying thickness and provides pathology with (another) unbiased tool which may help to reduce the interobserver variability among pathologists. We also show that a new class of powerful proteomic-technologies, here the Evosep-LC – timsTOF HT, provide fast and high repeatability. Crucially, we also show that specific protein content correlates with important histological lesion severity in kidney transplantation. We also believe that future research is needed to validate the method presented here for clinical use, which could significantly improve kidney transplant diagnosis and ultimately prognosis. In general, we see progress in this area. Through integration and establishment in the clinic, these new methods, which generate high-density information in the gigabyte range from FFPE samples within minutes, can open up great opportunities, including new insights in pathology and new possibilities for diagnosis.

## Author Contributions

JK and GC conceived and designed the study. RH, KM, RB and AC optimized sample preparation and analysis. RH and NC collected and prepared the patient cohort for analysis. KM and RB analyzed the cohort. RH, HPS and JK performed data analyses and visualizations. RH, HPS, JK and GC wrote the initial draft of the manuscript. RH created the figures. All authors subsequently read and revised the manuscript and read and approved the final version.

