## Supplementary data for "Highly Repeatable Tissue Proteomics for Kidney Transplant Pathology: Technical and Biological Validation of Protein Analysis using LC-MS/MS"

Supplementary table 1.
Inter quantile range (IQR) of coefficient of variance of technical and biological replicates

| Group | Condition | Lower | Median | Upper |
|---|---|---|---|---|
| **Technical Replicates** | 10 µm 1 | 5.8 | 10.4 | 18.6 |
| | 10 µm 2 | 4.9 | 9.5 | 17.2 |
| | 10 µm 3 | 5.5 | 10.3 | 18.2 |
| | 10 µm 4 | 6.0 | 11.4 | 20.5 |
| | 10 µm 5 | 5.2 | 10.2 | 18.7 |
| | 10 µm 6 | 5.4 | 10.2 | 17.6 |
| | 5 µm 1 | 7.9 | 15.6 | 28.5 |
| | 5 µm 2 | 8.4 | 15.5 | 29.6 |
| | 5 µm 3 | 10.3 | 20.7 | 38.2 |
| | 5 µm 4 | 13.8 | 27.1 | 53.5 |
| | 5 µm 5 | 11.5 | 21.7 | 41 |
| | 5 µm 6 | 15.4 | 27.0 | 62.9 |
| **Biological Replicates** | 5 µm | 12.6 | 19.4 | 30.7 |
| | 10 µm | 6.6 | 10.2 | 15.4 |

Supplementary figure 1. Schematic Sample Preparation (A) and Analysis (B)

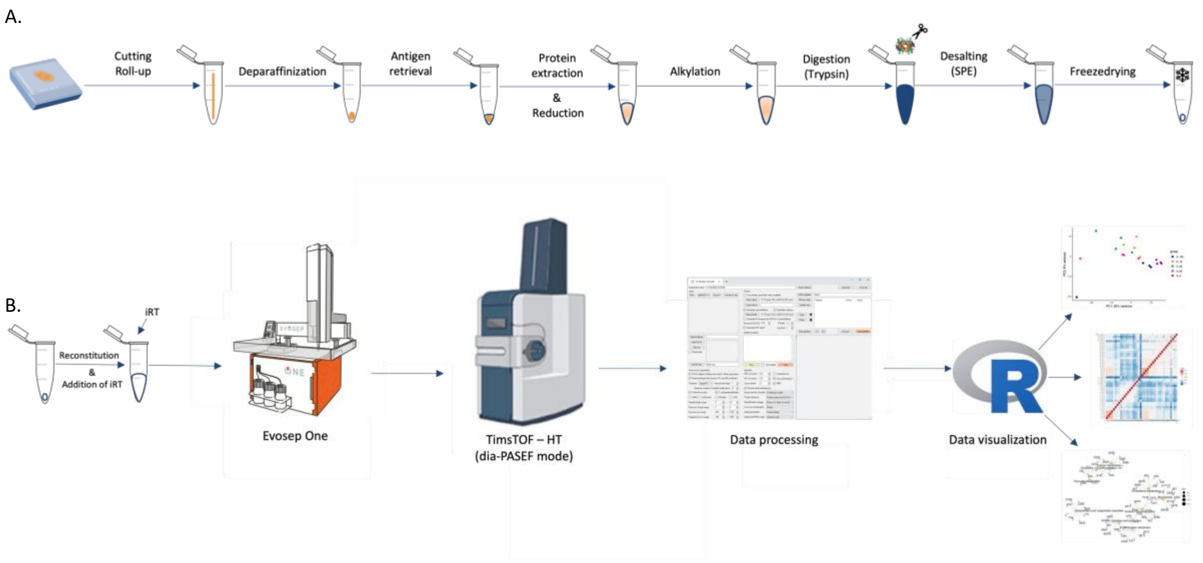
